# Comparative Transcriptome Analysis of Hypocotyls During the Developmental Transition of C_3_ cotyledons to C_4_ Leaves in *Halimocnemis mollissima* Bunge

**DOI:** 10.1101/2022.05.13.491777

**Authors:** Mahdis Zolfaghar, Twan Rutten, Mohammad Reza Ghaffari, Ali Mohammad Banaei-Moghaddam

**Author notes:** **Correspondence:** Mohammad Reza Ghaffari, (M.R.G), Ali Mohammad Banaei-Moghaddam, (A.M.B-M).

## Abstract

Identification of signaling pathways that control C_4_ photosynthesis development is essential for introducing the C_4_ pathway into C_3_ crops. Species with dual photosynthesis in their life cycle are interesting models to study such regulatory mechanisms. The species used here *Halimocnemis mollissima* Bunge, belonging to the Caroxyleae tribe, displays C_3_ photosynthesis in its cotyledons and a NAD-ME subtype of C_4_ photosynthesis in the First leaves (FLs) onwards. We explored the long-distance signaling pathways that are probably implicated in the shoot-root coordination associated with the manifestation of the C_4_ traits, including efficient resource usage by comparing the mRNA content of hypocotyls before and after the C_4_ first leave’s formation. Histological examination showed the presence of C_3_ anatomy in cotyledons and C_4_ anatomy in the FLs. Our transcriptome analyses verified the performance of the NAD-ME subtype of C_4_ in FLs and revealed differential transcript abundance of several potential mobile regulators and their associated receptors or transporters in two developmentally different hypocotyls of *H. mollissima* Bunge. These differentially expressed genes (DEGs) belong to diverse functional groups, including various transcription factor (TF) families, phytohormones metabolism, and signaling peptides, part of which could be related to hypocotyl development. Our findings support the higher nitrogen and water use efficiency associated with C_4_ photosynthetic and provide insights into the coordinated above- and under-ground tissue communication during the developmental transition of C_3_ to C_4_ photosynthesis in this species.

## Introduction

As an adaptation to the hot and dry environment, C_4_ photosynthesis is a carbon concentration mechanism that reduces the counteracting photorespiration by suppressing the oxygenase activity of the carboxylating enzyme RuBisCO (Ribulose-1,5-bisphosphate carboxylase/oxygenase). The leaves of C_4_ plants have a unique structure called Kranz anatomy, enabling them to spatially separate two phases of photosynthesis into the mesophyll and bundle sheath cells. The Kranz anatomy precedes evolving C_4_ biochemistry (McKown and Dengler 2007). A universal feature of C_4_ plants is their higher usage efficiency of resources such as nitrogen, water, and radiation (Vogan and Sage 2011; Fatima *et al*. 2018). Nitrogen and water are uptaken by roots and are largely controlled by the demands in the shoots. The indispensable shoot-root communication is coordinated by long-distance signaling molecules such as phytohormones, signaling peptides, and TFs (Ko and Helariutta 2017).

Generally, in the C_4_ pathway, the carbon fixation begins in the mesophyll, where carbon dioxide is converted into bicarbonate and then to four-carbon (C_4_) organic acids. The C_4_ compounds are transported to the bundle sheath and decarboxylated to release the CO_2_ at the site of RuBisCO for assimilation (Sage 2004; Gowik and Westhoff 2011). Based on the primary decarboxylating enzymes, NADP-ME and NAD-ME are two main biochemical subtypes of the C_4_ pathway (Rao and Dixon 2016). As a striking example of convergent evolution, the conversion of C_3_ into C_4_ photosynthesis has evolved independently in more than 60 lineages of angiosperms (Slewinski 2013). The polyphyletic origin of C_4_ syndrome implies that the transition of C_3_ into C_4_ is not as genetically complicated as inferred from the associated anatomical and biochemical features. It is suggested that the pre-existing C_3_ gene regulatory networks are recruited in the C_4_ pathway with relatively few modifications (Hibberd and Covshoff 2010; Reyna-Llorens and Hibberd 2017).

In addition to efforts distinguishing genes encoding C_4_ enzymes, other studies have tried to identify the genetic architecture and regulation of initiating the C_4_ metabolism (Gowik *et al*. 2011; Wang, Vlad and Langdale 2016; Cui, 2021). For this transcriptome profiles were compared of C_3_ and C_4_ tissues of close relative (C_3_ and C_4_ *Flaveria*) (Gowik *et al*. 2011) or distantly related (C_3_ rice and C_4_ maize) (Wang *et al*. 2014) species. Most of the omic analyses on C_4_ photosynthesis have been performed in C_4_ plants belonging to the NADP-ME subtype (Gowik *et al*. 2011; Lauterbach *et al*. 2017). However, to comprehend the nature and commonality of regulatory features of C_4_ syndrome it is essential to investigate both biochemical subtypes. The nature and mechanism of distant signaling molecules act in C_4_ plants to coordinate the above- and under-ground tissues to maintain their homeostasis and establish C_4_ features, including Kranz anatomy, nitrogen, and water use efficiency, remains largely obscure and has hampered the long-standing goal of engineering C_4_ metabolism into the C_3_ crops (Gerlich *et al*. 2018).

Recently, species such as *Haloxylon ammodendron* (Y. Li *et al*. 2015) and *Salsola soda* (Lauterbach, Billakurthi et al. 2017) have been found to display two photosynthetic mechanisms in their life cycle. As such they provide an excellent model for studying the molecular regulatory mechanisms underlying the coordination of roots and modified shoots during the C_3_ to C_4_ conversion while eliminating phylogenetic noises that hindered earlier models. Unlike the C_3_ pathway in cotyledons, which coincides with moderate temperature, the C_4_ pathway in these plants, which belongs to the NADP-ME subtype of C_4_ photosynthesis, outperforms at higher light and temperature conditions concurrent with the formation of the first leaf stage onwards (Lauterbach, Billakurthi et al. 2017).

*H. mollissima* Bunge used in the present study belongs to the Caroxyleae tribe and performs both C_3_ (in cotyledons) and C_4_ photosynthesis (NAD-ME subtype, in FLs onwards) (Akhani *et al*. 2009). To unravel the signaling pathways associated with the C_3_ to C_4_ transition in *H. mollissima* Bunge, we used RNA sequencing to generate transcriptome profiles of hypocotyls before and after forming the C_4_ FLs alongside the FLs. Genes associated with several phytohormone metabolisms, TFs, and signal peptides were differentially expressed in hypocotyls before and after forming C_4_ FLs. Our results indicated that, besides those associated with hypocotyl development, some of these genes show homology to known shoot-to-root and vice-versa signals and their receptors or transporters that are possibly implicated in establishing unique C_4_ traits, especially in the efficient consumption of resources such as nitrogen and water.

## Material and Methods

### Plant Cultivation and Sampling

Seeds of *H. mollissima* Bunge were collected at Mahdasht, Karaj, Iran (35°43^’35.4 “^N; 50°44^’^04.5^”^E) in November 2019 and stored at 4°C. After removal of the hard seed coats, seeds were disinfected with ethanol 75%, washed with water and germinated on the wet, sterilized filter paper in Petri dishes. After root formation, at 3 days after incubation (DAI), seedlings were transferred into the plastic pots (5 cm diameter, 7 cm height) filled with sand, perlite, and compost. *H. mollissima* Bunge Plants were grown in a greenhouse at the Institute of Biochemistry and Biophysics, University of Tehran, at 25-30°C, ~200 μmol m^−2^ s^−1^ illumination and 16/8-hours light/dark cycle.

For sampling, two series of *H. mollissima* Bunge were cultivated. In the first series, expanded cotyledons, hypocotyls, and roots (Series A) (Fig. 1(a)) were harvested before FLs were formed. In the second series, FLs, cotyledons, hypocotyls, and roots (Series B) (Fig. 1(b)) were harvested after formation of FLs. Tissues of interest were sampled in January and February 2020 between 11:00 and 13:00, shock-frozen in liquid nitrogen and stored at −80°C. Two biological replicates of hypocotyls and FLs and three of all sampled tissues were used for RNA sequencing and RT-qPCR, respectively, each containing pooled tissue from 5 independent plants.

**Fig. 1.**
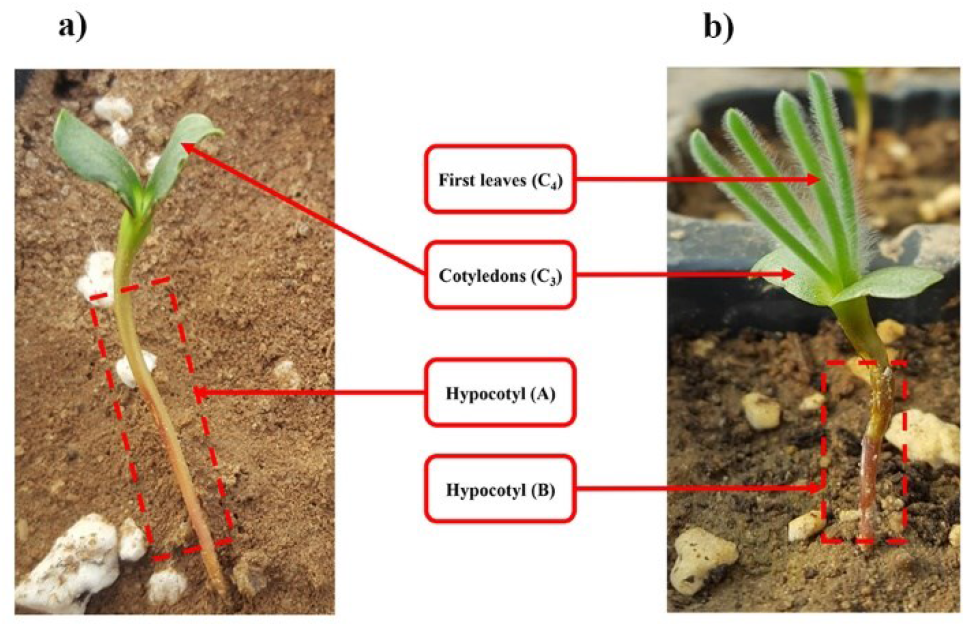
Sampled tissues from *H. mollissima* Bunge in the current study a) Series A b) Series B

### Light Microscopy

For histological studies, C_3_ cotyledon and C_4_ first leaf were fixated for 16h in 50 mM Phosphate Puffer, pH 7.0, containing 1 % (v/v) glutaraldehyde and 4 % (v/v) formaldehyde at 8°C. After 2 x 10 min washing with distilled water and subsequent dehydration in an ascending ethanol series (30, 40, 50, 60, 75, 90 % and 2 × 100 % ethanol, 10 min each), probes were infiltrated with Spurr resin and polymerized in an oven at 70°C for 24 h. Semithin sections of the middle regions of fully expanded cotyledons and FLs were made on a Reichert Ultracut S (Leica Microsystems, Wetzlar, Germany) and stained with 1 % (w/v) methylene blue and 1 % (w/v) Azur II. Images were taken on a Zeiss Axiovert 135 microscope (Carl Zeiss, Jena, Germany) and stored as TIFF files.

### RNA Extraction and Sequencing

The guanidine/ phenol-based method (Chomczynski and Sacchi 1987) was used to isolate total RNA from 100-150 mg of the homogenized samples. Quality, quantity, and integrity of isolated RNA were checked by Agarose gel electrophoresis, Nanodrop (Thermo Fisher Scientific), and Agilent Bioanalyzer 2100 system (Agilent Technologies Co. Ltd., Beijing, China), respectively. High-quality RNA (RIN value > 5.2) was used for cDNA preparation and sequencing by Illumina Hiseq 2500 platform at the Novogene Bioinformatics Institute (Beijing, China). One hundred fifty bp paired-end reads were obtained, and their adapter-contained and low-quality (Qscore< = 5) base-contained were removed.

### De novo Assembly and Functional Annotation

Raw reads were evaluated for quality by FastQC tool (http://www.bioinformatics.babraham.ac.uk/projects/fastqc/). Due to removing adapters and low-quality reads by the Novogene Bioinformatics Institute, there was no need for trimming. High-quality reads were assembled by Trinity v2.4.0 (Grabherr *et al*. 2011) using default settings. First, probable open reading frames (ORF) were predicted using the Transdecoder tool (Haas and Papanicolaou, 2016) with a minimum length of 100 amino acids. To obtain Gene Ontology (GO) terms of the transcripts, Blast2GO software was used with 1e^3^ as an e-value cut-off to perform BlastX against a non-redundant protein database. The online KEGG Automatic Annotation Server (KAAS) (http://www.genome.jp/kegg/kaas) and Mapman V4.0 (https://www.plabipd.de/portal/mercator4) (Schwacke *et al*. 2019) were used for pathway analysis (p-value ≤ 0.05). Also, the Arabidopsis TAIR locus IDs were assigned to each transcript during Mapman analysis (https://www.plabipd.de/portal/web/guest/mercator-sequence-annotation) (Lohse *et al*. 2014).

### Determination of Genes with Different Expression

RSEM tool (http://deweylab.biostat.wisc.edu/RSEM) (Li and Dewey 2011) was used to estimate the abundance of the transcripts for each sample. The high-quality raw reads were separately mapped back onto the assembled transcripts using Bowtie 2.0. Differential expression analysis was performed by edgeR (Robinson, McCarthy and Smyth 2010) in R after normalization using the trimmed mean of M-values (TMM) method. Statistical tests for the pairwise comparisons were Benjamini-Hochberg correction FDR ≤ 0.05, log_2_FC ≥ 1.5 or ≤ −1.5 to set a significant threshold for determining DEGs. Principal component analysis (PCA) with log_2_ transformed read count (Transcript Per Million TPM) was performed in RStudio to visualize the expression pattern of the samples. Furthermore, hierarchical clustering of the transcript abundance of selected known C_4_ proteins using Pearson correlation, average linkage method was performed (http://www.heatmapper.ca/expression/).

The DEGs most likely associated with the hypocotyl development of *H. mollissima* Bunge were excluded from further analysis by comparing them with DEGs involved in the hypocotyl development in other plants of flax and hemp (Roach and Deyholos 2008; Behr *et al*. 2018) (https://bioinformatics.psb.ugent.be/webtools/Venn/).

### Determination of Active Biological Functions

Pathways in each tissue of interest were explored using the percentage of total transcripts (normalized to Transcript Per Million (TPM)) of each particular bincode in Mapman V4.0. According to the further emphasis of this study on the survey of probable long-distance signaling, gene enrichment analysis was conducted by PageMan software (Usadel *et al*. 2006) through Fisher’s exact test followed by the Benjamini-Hochberg correction to identify significant differences of Mapman bins among the two developmentally different hypocotyls. Moreover, Mapman V3.6.0RC1 was used to visualize selected DEGs (through the generation of a custom MapMan pathway image) encoding phytohormone synthesizing and degrading enzymes, transporter (“Phytohormone action bin” and “Solute transporter”), and also genes related to different signal transduction pathways (“Multi-process regulation. Calcium-dependent signaling”, “Protein modification. Protein kinase” and “Redox homeostasis” bins).

### Identification of TFs and Prediction of Cis-acting elements in the Promoter of Predicted Mobile TFs

To predict differentially expressed TFs between hypocotyl before and after the formation of the FLs, the homology of related amino acid sequences against the plant TFs database (PlantTFDB V5.0, http://planttfdb.gao-lab.org/prediction.php) was implemented. Afterward, TFs with mobile mRNA, based on the information about the genes with “cell-to-cell mobile mRNA” in The Arabidopsis Information Resource (TAIR) database, were predicted.

Since the *H. mollissima* Bunge genome is not yet available, the promoter region of the orthologs of predicted mobile TFs in Arabidopsis was obtained from the Arabidopsis Gene Regulatory Information Server (AGRIS) database (https://agris-knowledgebase.org/AtcisDB/), and their cis-acting were predicted using PlantCARE (http://bioinformatics.psb.ugent.be/webtools/plantcare/html/).

### Real-time quantitative PCR (RT-qPCR)

To confirm RNA seq data, RT-qPCR was performed for eight genes. Total RNA was extracted by the above-mentioned method (Chomczynski and Sacchi, 1987) and treated with DNase I (Thermo Fisher). RNA was reverse transcribed with the AddScript cDNA Synthesis Kit (add bio), according to manufacturer’s instructions, using 1 μg of RNA. RT-qPCR was performed with a 6plex real-time PCR system (Qiagen, Rotor Gen Q, Germany) using the RealQ Plus 2x Master Mix Green kit, without ROX (AMPLIQON) using 1 μL of 1:40-fold diluted template cDNAs. Run conditions were: 10 min of initial denaturation at 95°C, 95°C for 30s, 60°C for 30s, and 72°C for 30s (40 cycles). Melting curve analysis and agarose gel electrophoresis were done to verify the primer specificity. Primer efficiency was estimated for each primer pair by four consecutive 10-fold dilutions of the cDNAs as a template. The relative expression of genes was calculated using the 2^−ΔΔCt^ method (Livak and Schmittgen 2001) using the Polyubiquitin ortholog from *H. mollissima* Bunge as an endogenous control. Three biological replicates were quantified for each sample.

## Results

### Cotyledons and FLs of *H. mollissima* Bunge have Different Anatomical Features in Accordance with Their Photosynthesis Strategy

C_3_ cotyledons of *H. mollissima* Bunge are flattened, bifacial, and their cross-section (Fig. 2 (a)) doesn’t show any Kranz anatomy. They have several layers of chlorenchyma cells around the vascular bundles (VBs) and intercellular airspaces across the blades (Fig. 2 (a)). While C_4_ first leaf is terete, linear, and covered by hairs, and its anatomy is salsoloid. The leaf cross section (Fig. 2 (b) and (c)) shows a large central water storage tissue which is respectively surrounded by the Kranz layer, palisade mesophyll cells, and a single-layered epidermis, while there is no space to be seen in the microscopic leaf structure.

**Fig. 2.**
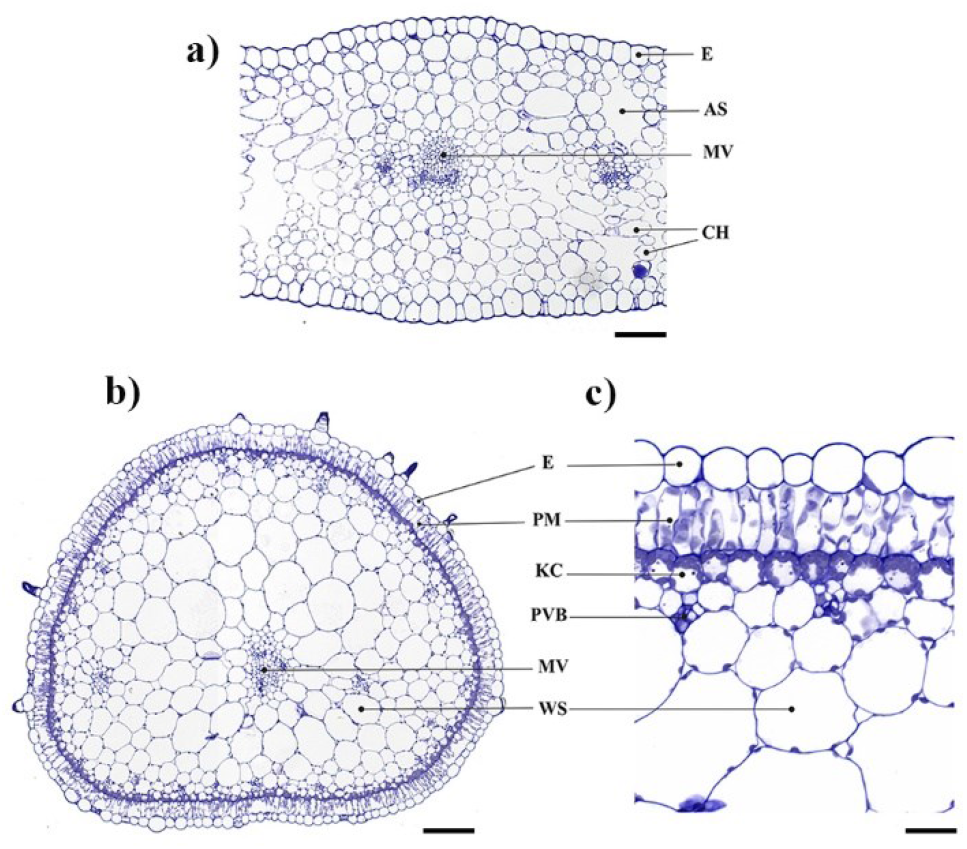
Leaf and cotyledon cross-sections of *H. mollissima* Bunge a) Cross-section of the cotyledon (C_3_), b) Cross-section of the entire leaf (C_4_, Salsoloid type Kranz anatomy), and c) close-up of the leaf Kranz anatomy Abbreviations: E, Epidermis; PM, palisade mesophyll; KC, Kranz cell; PVB, peripheral vascular bundle, MV, mid-vein, WS, water-storage tissue; AS, air space; CH, chlorenchyma. Bars = 100 μm (a, b), 25 μm (c)

### Processing of Raw Reads and Statistics of De novo Assembly

The mRNA of three different tissues, including hypocotyls before (hypocotyl A) and after (hypocotyl B) the formation of FLs, and the FLs themselves (FL), were sequenced. The quality of reads was checked, and based on FastQC results (Phred score= 36) all reads had high quality (Additional file 1(A)) circumventing the need for further trimming. Raw data is available in the ? database under accession number ? (Additional file 2). To evaluate the Trinity results, N50 and total assembly length were checked by Trinitystat.pl (Additional file 1 (B)). PCA analysis (Fig. 3), where the first principal component contained 69.57% and the second principal component had 21.52% of the variance, demonstrated that the analyzed tissues have a distinct expression patterns. Nevertheless, the replicates’ expression data were similar and grouped. As expected, the same tissues (hypocotyl A and hypocotyl B) had more similar expression patterns than the FLs.

**Fig. 3.**
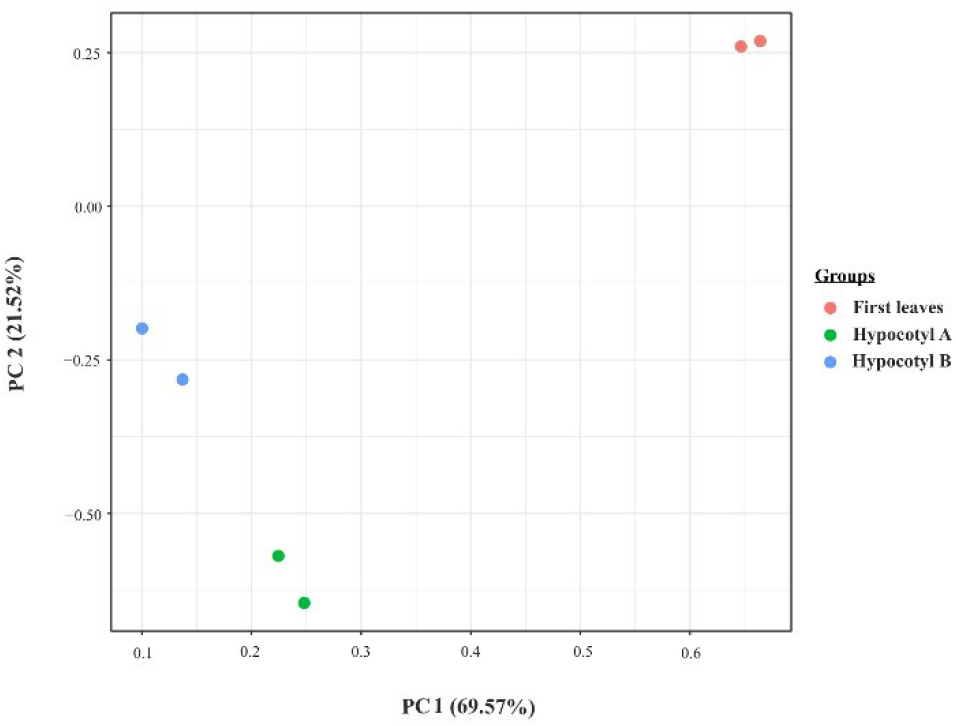
PCA analysis which explains 91.09% of the total variation. Principal component analysis of the read counts separated the samples based on the tissue type and indicated the consistency and validation of our normalized data

### Determination of Active Functional Pathways in analyzed tissues

To evaluate how the transcriptome as whole changes during the transition of photosynthesis strategy, the functional classes of each tissue transcriptome were determined as the percentage of all transcripts (normalized to Transcript Per Million (TPM)) assigned to each Mapman bincodes (Fig. 4(a), Additional file 3). MapMan analysis showed “Not assigned, annotated” and “Not assigned, not annotated” with 19.41 and 45.86% of all transcripts being the most significant functional gene categories in each sample. While with 15.44% the “Photosynthesis” bin was the third-highest abundant functional class in the FLs, in the hypocotyl A and B it accounted for only 3.25% and 2.47%, respectively. Out of 16 transcripts associated with known C_4_ proteins, 14 transcripts, including Beta-carbonic anhydrase (BCA3), PEP carboxylase1 (PEPC1), pyruvate orthophosphate dikinase (PPdK), aspartate aminotransferase (Asp-AT), alanine aminotransferase (Ala-AT), NAD-dependent malic enzyme (NAD-ME), NADP-dependent malic enzyme (NADP-ME), Triosephosphate translocator (TPT), Pyrophosphatase (PPase6), pyruvate orthophosphate dikinase related protein1 (PPdK-RP1), PEP/phosphate translocator (PPT2), Bile acid:sodium symporter family protein (BASS4), Bile acid:sodium symporter family protein (BASS2), Adenosine monophosphate kinase (AMK2) were more abundant in FLs than hypocotyls (Fig. 4(b), Additional file 4). As expected, NAD-ME is more abundant than NADP-ME in FLs.

**Fig. 4.**
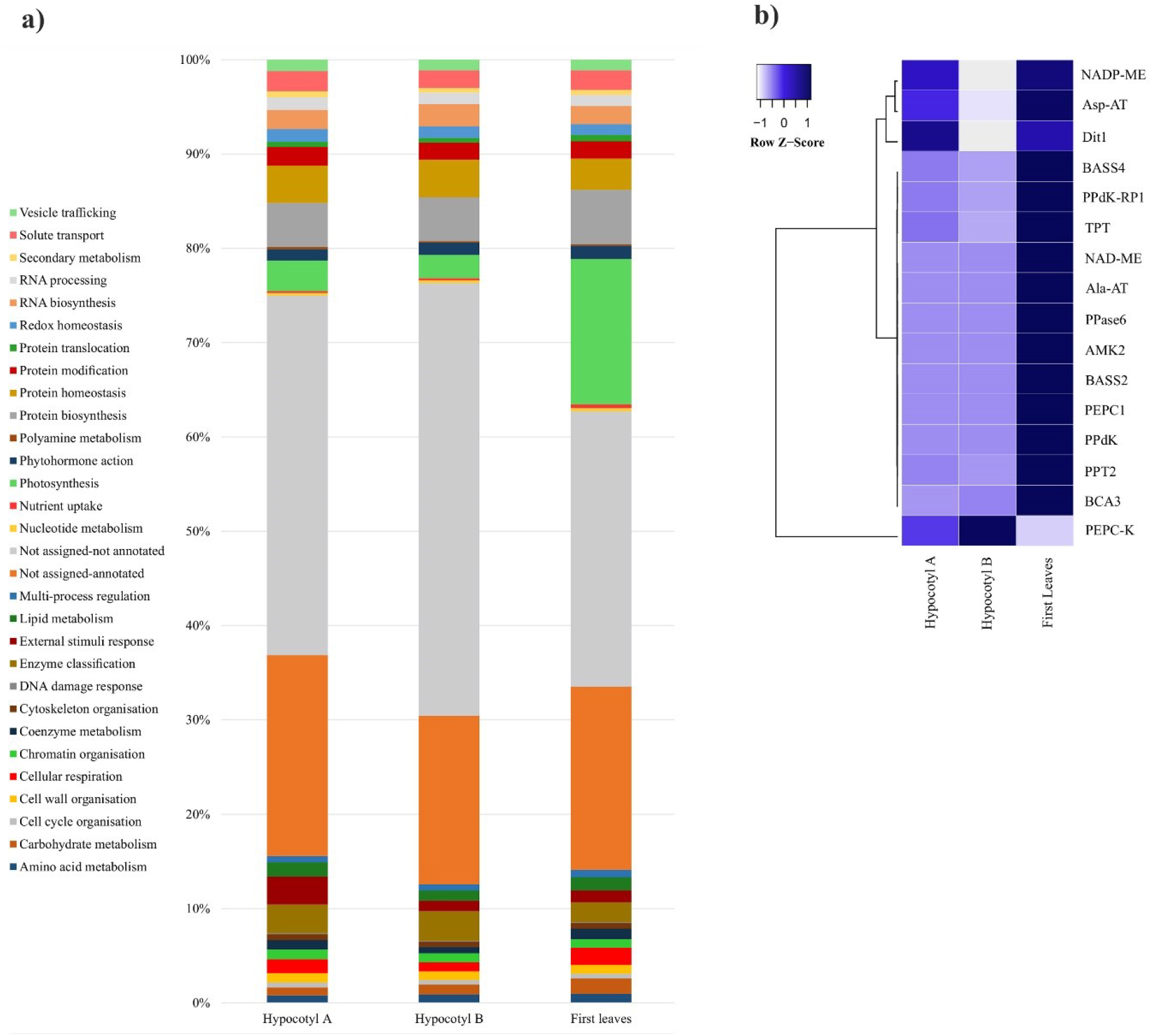
a) Active functional classes of each sample according to all percentages of all transcripts belonged to Mapman categories **b)** Relative transcript abundance of C_4_ cycle proteins: most C_4_ proteins are more expressed in leaves than in hypocotyls. Beta-carbonic anhydrase (BCA3), PEP carboxylase1 (PEPC1), pyruvate orthophosphate dikinase (PPdK), aspartate aminotransferase (Asp-AT), alanine aminotransferase (Ala-AT), NAD-dependent malic enzyme (NAD-ME), NADP-dependent malic enzyme (NADP-ME), Triosephosphate translocator (TPT), Pyrophosphatase (PPase6), pyruvate orthophosphate dikinase related protein1 (PPdK-RP1), PEP/phosphate translocator (PPT2), Bile acid:sodium symporter family protein (BASS4), Bile acid:sodium symporter family protein (BASS2), Adenosine monophosphate kinase (AMK2), Dicarboxylate translocator1 (Dit1), phosphoenolpyruvate carboxykinase (PEPC-K)

“Protein biosynthesis” and “Protein homeostasis” were the third- and fourth-highest abundant functional categories in hypocotyl A (4.68 and 3.96%, respectively) and B (4.67 and 4%, respectively) (Fig. 4(a)). “RNA biosynthesis” (1.92-2.34%) and “solute transport” (1.87-2.09%) were also abundant in all three samples (Additional file 3). Although the abundance of most functional categories is similar in hypocotyl A and B, the “RNA biosynthesis” (2.04 and 2.34 %, respectively) and “Phytohormone action” (1.2 and 1.31%, respectively) classes are somewhat more abundant in the hypocotyl B.

The Pairwise comparison between the samples revealed several DEGs (FDR<= 0.05) among three different developmental tissues (Additional file 5). In order to identify possible signaling pathways involved in the C_3_ to C_4_ transition, pathways enrichment analysis was conducted for 8424 DEGs in the hypocotyl B vs. hypocotyl A comparison. Among the pathways that according to the overrepresentation analysis appeared to be up-regulatedly enriched in hypocotyl B were those involved in “Protein biosynthesis”, “Protein homeostasis and the ubiquitin-proteasome system” (Fig. 5).

**Fig. 5.**
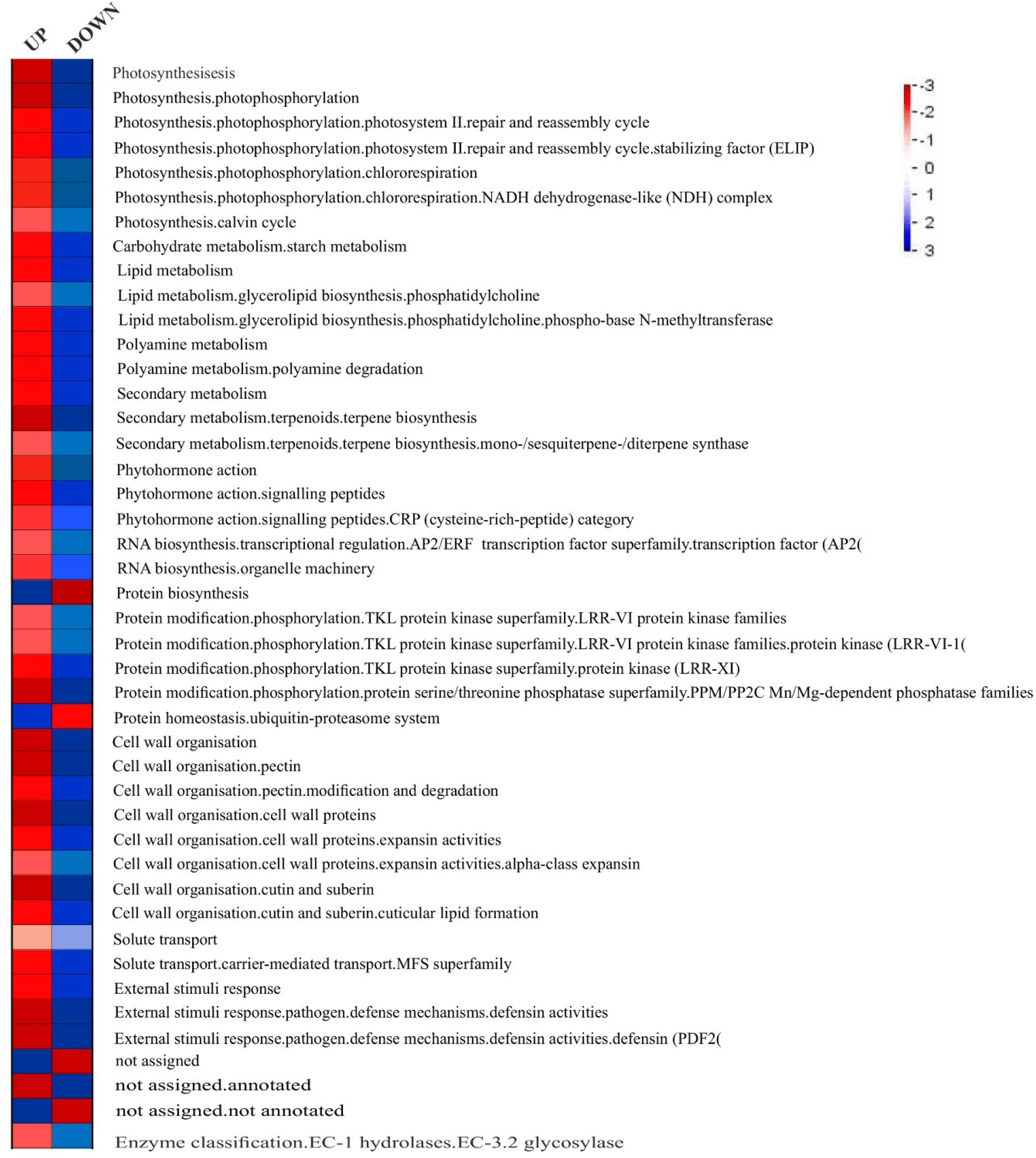
Overrepresentation analysis of up- and down-regulated genes in pairwise comparison of hypocotyl B vs. hypocotyl A within functional gene classes defined by MapMan bins (MapMan, v3.6, available online at https://www.plabipd.de/portal/mercator4) in Pageman software. Fisher’s exact test, followed by the Benjamini Hochberg correction and cut-off value one, was used for functional enrichment of DEGs between three binary comparisons Blue, up- or downregulated genes are significantly overrepresented; red, up- or downregulated genes are significantly underrepresented

Next, to explore possible long-distance regulators, we focus on the various regulatory mechanisms by further analyzing DEGs in the two developmentally different hypocotyls. To differentiate between genes controlling the transition of photosynthesis strategy and those involved in hypocotyl development, we compared them (based on the assigned Arabidopsis IDs) with previously identified genes associated with hypocotyls development (Roach and Deyholos, 2008; Behr *et al*. 2018). This way 190 genes were excluded (Additional file 6, Additional file 7). We then focused on the unique genes which do not overlap with related developmental genes.

### DEGs Associated with Classical Phytohormones Metabolism

Mapping of unique DEGs of hypocotyl B vs. hypocotyl A in Mapman V3.6.0RC1 (Fig. 6, Additional file 8) indicated several genes involved in the biosynthesis, degradation, transport, and signal transduction of plant hormones (Phytohormone bin in Mapman) that are up- or down-regulated after the establishment of C_4_ photosynthesis in the FLs.

**Fig. 6.**
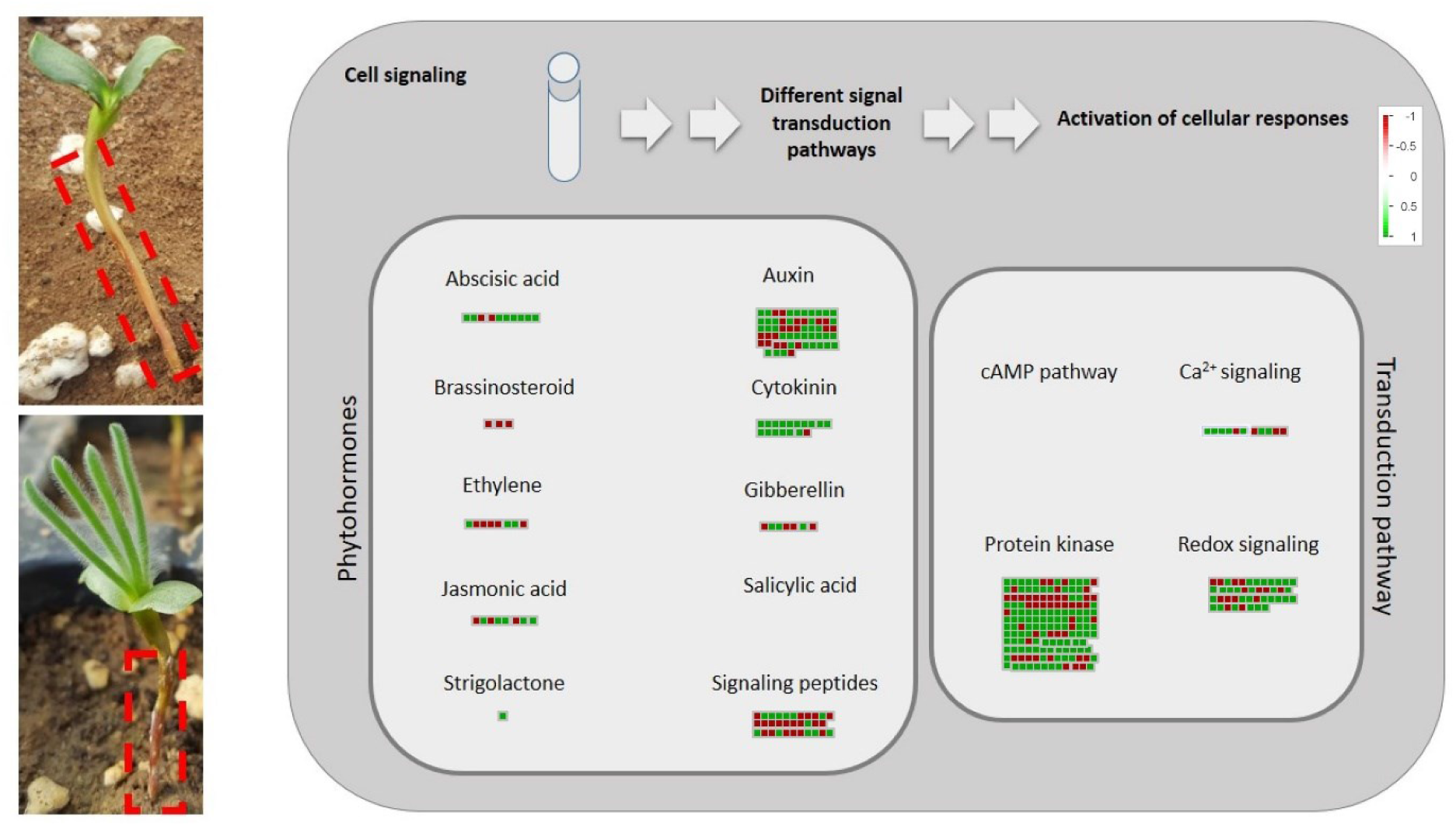
“Cell signaling” using MapMan. Visualization of transcriptional changes in genes involved in the various signaling pathways (FDR<0.05, log2FC>1.5 <−1.5) at hypocotyl B Vs. hypocotyl A (DEGs encoding phytohormone synthesizing and degrading enzymes, transporter (“Phytohormone action bin” and “Solute transporter”), and different signal transduction pathways (“Multi-process regulation. Calcium-dependent signalling", “Protein modification. Protein kinase” and “Redox hemoestasis” bins));Red, down-regulated gene, and green, up-regulated genes

Several transcripts encoding membrane bound auxin transporters, such as PIN-FORMED (PIN) (PIN2; At5g57090), and membrane-bound ATP-binding cassette-B (ABCB) auxin transporter (ABCB19; At3g28860), which are involved in long-distance auxin signaling proved to be less abundant in hypocotyl B compared to hypocotyl A. Other auxin-responsive genes like NITRATE TRANSPORTER 1 (NRT1; AT1G12110), an auxin-responsive transporter, were down-regulated in hypocotyl B after C_4_ establishment, as was the flavin-dependent monooxygenase (YUCCA) involved in auxin biosynthesis. In contrast, transcripts of several PIN-Like transporters (PILS), which are distributed in the intracellular compartments and mediate intracellular auxin homeostasis, were more abundant in hypocotyl B. Also transcripts assigned to AUX/ IAA (AT5G43700, AT3G04730, and AT4G29080) and ARF7/ARF19 (AT1G19220) auxin signal transduction genes were up-regulated in hypocotyl B. Apparently, the biosynthesis and long-distance transport of auxin are reduced in hypocotyls after the formation of the FLs. In contrast to this, almost all transcripts associated with different cytokinin pathways bin (biosynthesis, signaling and transport) were up-regulated in hypocotyl B (Fig. 6). For example, type-B ARABIDOPSIS RESPONSE REGULATOR (ARR) and type-A ARR, involved in the cytokinin signaling, were up- and down-regulated in the hypocotyl B, respectively. Also, transcripts of ENT (EQUILIBRATIVE NUCLEOTIDE TRANSPORTER 1) (at4g05120, at1g70330), importers to take up apoplastic nucleosides of cytokinins, were more expressed in the hypocotyl B. DEGs associated with brassinosteroids were down-regulated.

Several long-range hormone transporters were more expressed in hypocotyl B (Fig. 6, Additional file 8), such as ABCG30 (AT4G15230), ABCG40 (AT1G15520), LHT1 (LYSINE-HISTIDINE TRANSPORTER) (at5g40780). NPF3.1 (Nitrate transport1/Peptide transporter family) (at1g68570), which is a multihormone transporter involved in the GA/ABA antagonism (Hirner *et al*. 2006), was down-regulated in hypocotyl B.

### Differentially Expressed Signaling Peptides

Overrepresentation analysis of DEGs in hypocotyls before and after the formation of the FLs revealed that in contrast to the CRP classes of plant peptides (showing an overrepresentation of down-regulated genes), transcripts related to NCRP did not show enrichment in hypocotyls after the formation of the FLs (Fig. 5). Transcript abundance of different groups of these classes, based on Mapman bins, was then compared between analyzed tissues which showed a complex pattern of up and down-regulation (Fig. 7, Additional file 4). In general, transcripts belonging to different groups of the CRP categories were less abundant in hypocotyls after the establishment of C_4_ photosynthesis in the FLs (Fig. 7 (a)). In this regard, transcripts assigned to several GASA group members, including GASA1 (at1g74670), GASA8 (AT2G39540), GASA13 (AT3G10185), GASA14 (AT5G14920), and GASA 10 (AT5G59845) as well as RALF signaling peptides like RALF33 (AT4G15800) and its receptor CrRLK1L were down-regulated in hypocotyl after C_4_ manifestation. Transcripts encoding peptides belonging to NCRP classes, including plant natriuretic peptide (PNP), CLAVATA3-EMBRYO-SURROUNDING REGION (CLE), Casparian strip integrity factor (CIF), PAMP-INDUCED SECRETED PEPTIDE (PIP) were more abundant in the hypocotyl after the formation of FLs (Fig. 6, Fig. 7 (b)). However, transcripts that are assigned to the receptor of C-terminally Encoded Peptides (CEP) such as CEPR2 (AT1G72180), acting in the nitrogen uptake signaling and Root meristem growth factor, were down-regulated in the hypocotyl after FLs formation.

**Fig. 7.**
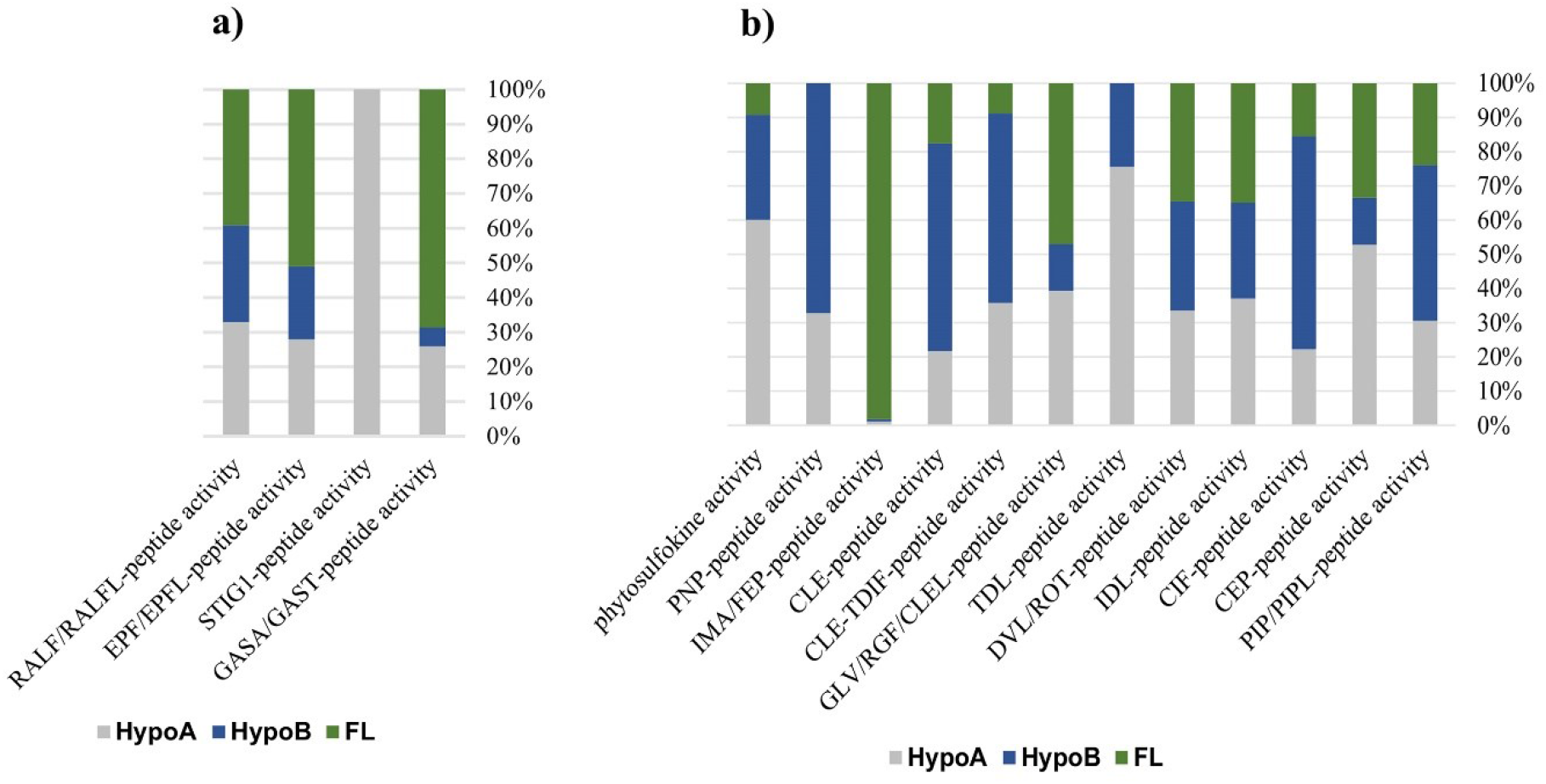
Relative transcript abundance of transcripts associated with a) CRP peptide activities b) NCRP peptide activities

### DEGs Taken Part in Various Signal Transduction Pathways

The binding of the ligands to their receptors at the cell surface triggers intracellular reaction cascades to amplify the input signals, eventually altering gene expression in various pathways. We found that transcripts of genes involved in the transduction pathways were differentially regulated in the hypocotyls after forming the FLs (Fig. 6). Transcripts of several protein kinase genes belonging to the casein kinase (CK) and sterility (STE) protein kinase superfamilies, which are implicated in responses to many signals, were more abundant in hypocotyl after forming the FLs. Most genes belonging to the AGC (PKA-PKG-PKC), and CMGC (Cyclin-dependent kinases, Mitogen-activated protein kinases (MAPK), Glycogen synthase kinases, and Cyclin-dependent like kinases) superfamilies were more expressed in hypocotyl B, as were transcripts associated with receptor-like cytoplasmic protein kinases (RLCKs).

Also several genes involved in redox signaling and also those encoding for calmodulin-like proteins (CMLs) involved in plants Ca2+ signaling processes were up-regulated in the hypocotyl after the formation of the FLs (Fig. 6).

### Differentially Expressed TFs

According to PlantTFDB V5.0, 290 out of 9640 ORFs from differentially expressed transcripts between hypocotyl before and after the formation of the FLs were annotated as different TF families with the best fit in *A. thaliana* locus IDs (Additional file 9). The mRNA levels of 71 and 219 of these TFs were down- and up-regulated, respectively, in hypocotyl after the formation of the FLs. In this comparison, C2H2, bHLH, and NAC families had a larger number of differentially regulated transcripts. Almost all transcripts belonging to C2H2, C3H, GRAS, MYB, TCP, Trihelix, NF-Y, and WRKY TFs had more expression in hypocotyl after the formation of the FLs. SHORT ROOT (SHR) (AT4G37650), a member of the GRAS family, and ELONGATED HYPOCOTYL 5 (HY5) (AT5G11260, AT3G17609), a member of the basic leucine zipper (bZIP) family, were up-regulated in hypocotyl after the formation of the FLs. Notably, both SHR and HY5 are shown to have mobility and could affect the expression of other genes (Dolan 2001; Chen *et al*. 2016).

Among 290 differentially expressed transcripts that were annotated as TFs based on PlantTFDB, 12 of them had orthologous with cell-to-cell mobile mRNA, according to the TAIR database (Table 1). Eight of these TFs were more abundant in hypocotyl after the formation of the FLs, including SCL1 (GRAS family, AT1G21450), HAT22 (HD-ZIP family, AT4G37790), HSFA1E (HSF family, AT3G02990), CDC5 (MYB_related, AT1G09770), MYBH (MYB_related family, AT5G47390), NAC053 (NAC family, AT3G10500), NAP (NAC family, AT1G69490) and ATWRKY40 (WRKY family, AT1G808_4_0).

**Table 1.**
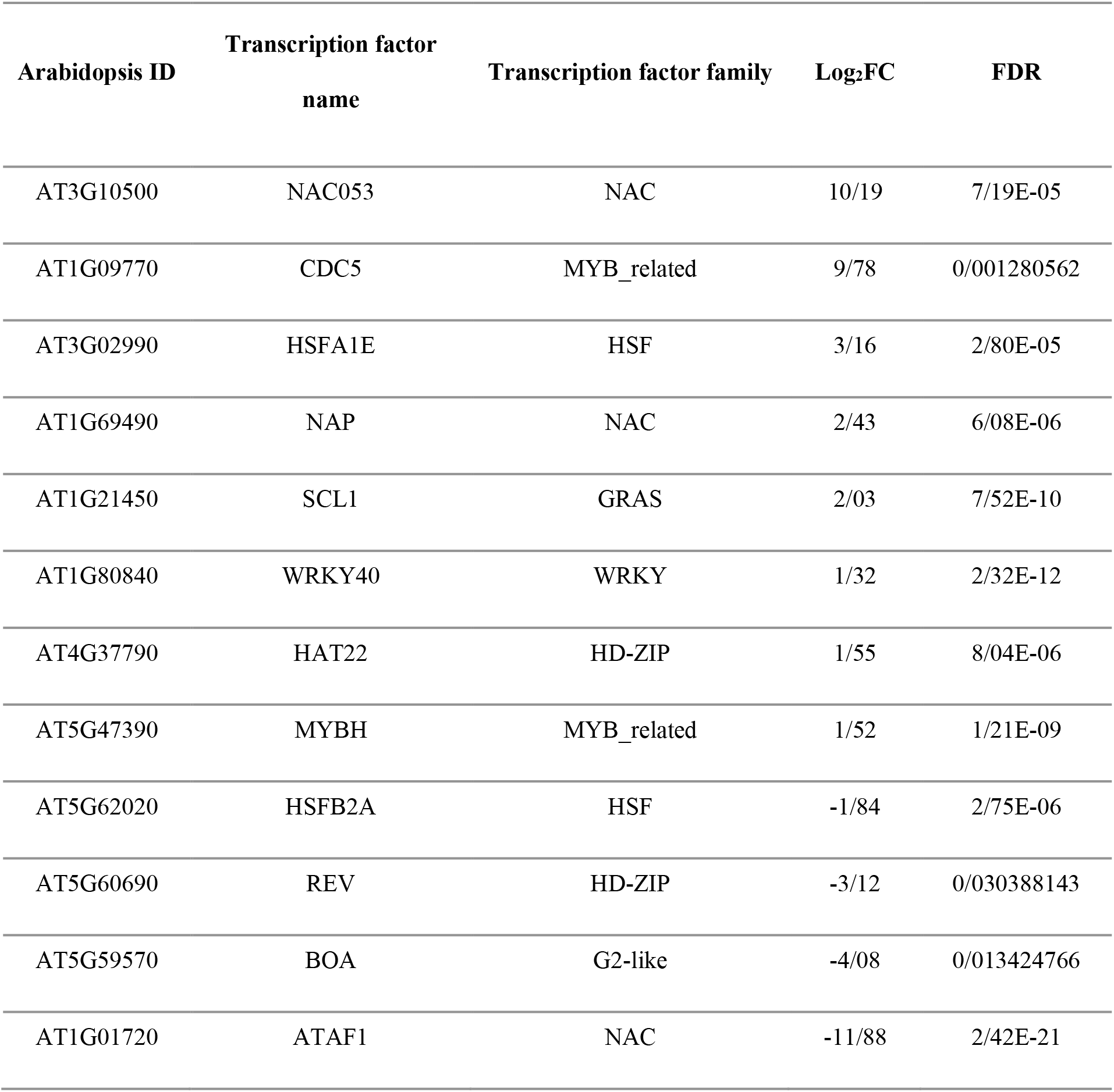
List of differentially expressed transcription factors between hypocotyl A and hypocotyl B with mobile mRNAs based on TAIR database (https://www.arabidopsis.org/)

| Arabidopsis ID | Transcription factor<br>name | Transcription factor family | Log<sub>2</sub>FC | FDR |
| --- | --- | --- | --- | --- |
| AT3G10500 | NAC053 | NAC | 10/19 | 7/19E-05 |
| AT1G09770 | CDC5 | MYB_related | 9/78 | 0/001280562 |
| AT3G02990 | HSFA1E | HSF | 3/16 | 2/80E-05 |
| AT1G69490 | NAP | NAC | 2/43 | 6/08E-06 |
| AT1G21450 | SCL1 | GRAS | 2/03 | 7/52E-10 |
| AT1G80840 | WRKY40 | WRKY | 1/32 | 2/32E-12 |
| AT4G37790 | HAT22 | HD-ZIP | 1/55 | 8/04E-06 |
| AT5G47390 | MYBH | MYB_related | 1/52 | 1/21E-09 |
| AT5G62020 | HSFB2A | HSF | -1/84 | 2/75E-06 |
| AT5G60690 | REV | HD-ZIP | -3/12 | 0/030388143 |
| AT5G59570 | BOA | G2-like | -4/08 | 0/013424766 |
| AT1G01720 | ATAF1 | NAC | -11/88 | 2/42E-21 |

To predict if specific cis-regulatory elements are overrepresented in the regulatory region of candidate mobile TFs with differentially expression levels in hypocotyls, PlantCARE (Lescot *et al*. 2002) was used to analyze the upstream promoter elements of the Arabidopsis orthologs of these transcripts. The results showed that Arabidopsis orthologs of differentially regulated mobile TFs carry several different cis-elements responsive to phytohormones and environmental stimuli such as light and temperature. Among these, Box 4 (part of a conserved DNA module involved in light responsiveness), AE-box (part of a module for light response), G-Box (cis-acting regulatory element involved in light responsiveness), and GT1-motif (light-responsive element) were more common light-responsive elements in the upstream of the mobile TFs (Additional file 10). Besides, the ABA-responsive element (ABRE) was predicted in the promoter of BOA, SCL1, HAT22, MYBH, NAC053, ATAF1, NAP, and WRKY40 TFs. Other motifs related to stress responses, including the MYB binding site involved in drought-inducibility (MBS), a cis-acting element involved in defense and stress responsiveness (TC-rich repeat), and a cis-acting regulatory element involved in the MeJA-responsiveness (CGTCA-motif and TGACG-motif), were predicted to be present in the upstream region of the orthologs of differentially regulated mobile TFs genes after the formation of the FLs.

To confirm RNA sequencing results, RT-qPCR was performed for eight target genes. The results were correlated with the RNA seq analysis (R^2^=0.81; Additional file 11). The RT-qPCR data analysis confirmed a higher abundance of the C_4_ enzymes, PEPC, BCA, and NAD-ME encoding genes in the FLs compared to the cotyledons (Fig. 8(a)). The expression pattern of the analyzed genes somewhat varied between developmentally different cotyledons and hypocotyls (Fig. 8(b), 8(d)). PEPC, BCA, NAD-ME, CEPR2, AUX/ IAA, ABCG40, WRKY40, and NRT1.1 were less expressed in the roots after the formation of the FLs (Fig. 8(c)).

**Fig. 8.**
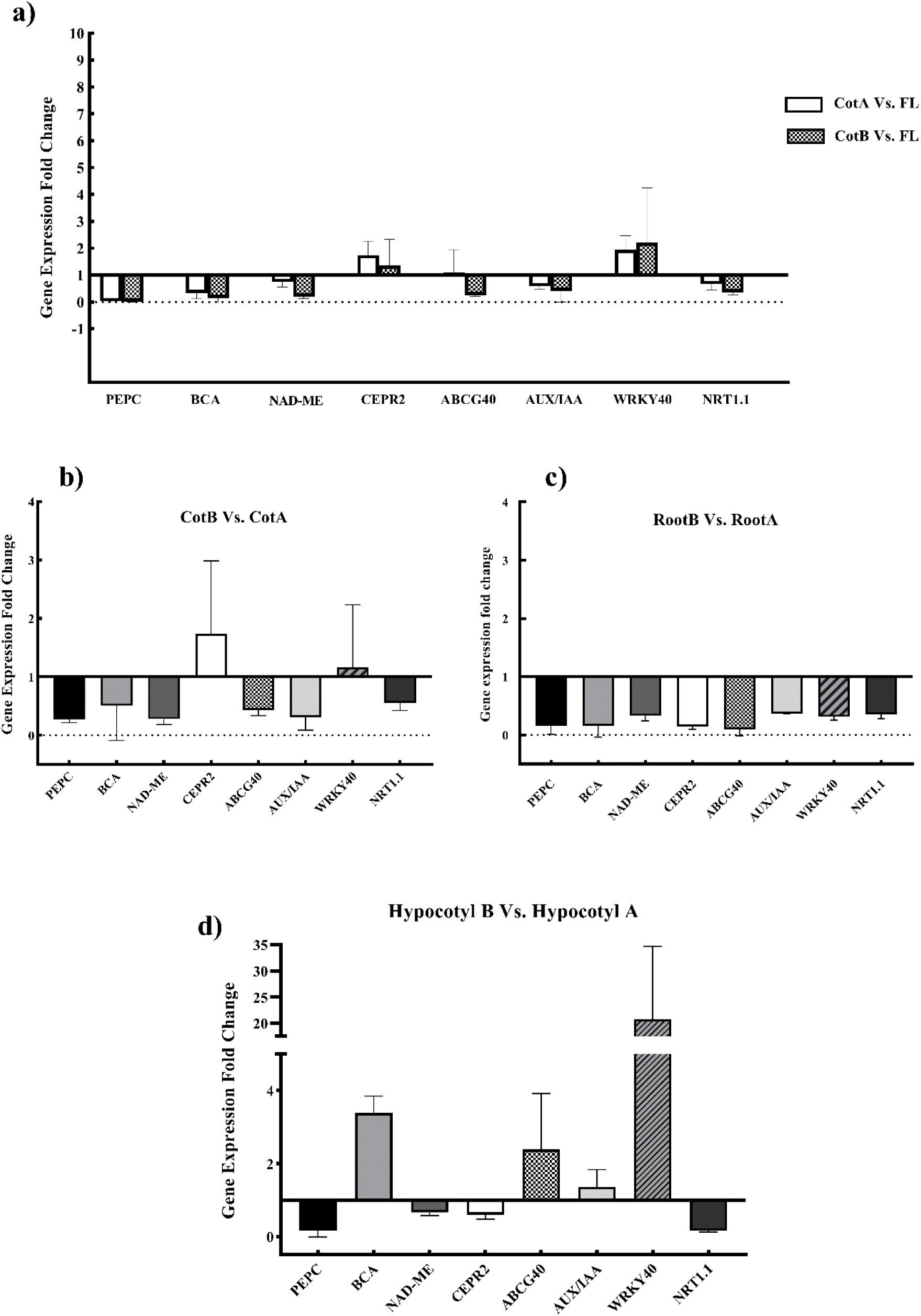
RT-qPCR analysis of selected genes in the tissues of interest a) (CotA, CotB Vs, FL) b) CotB Vs. CotA c) RootB Vs. RootA d) Hypocotyl BVs. Hypocotyl A

## Discussion

### Species with Switching Mechanisms are Valuable Models for studying C_4_ Photosynthesis

We investigated the transcriptome profile of *H. mollissima* Bunge hypocotyls before and after first leaf formation to identify possible components of long-distance signaling pathways involved in the unique C_4_ traits through the connection between the above- and under-ground tissues. *H. mollissima* Bunge belongs to the Caroxyleae tribe and has two photosynthetic mechanisms in its annual life cycle; C_3_ in cotyledons and NAD-ME subtype of C_4_ in the FLs onwards. Anatomical studies of cotyledons and FLs (Fig. 2) reveal features related to C_3_ and C_4_ (Salsoloid) (Akhani *et al*. 2009; Freitag and Kadereit 2014), respectively. In leaves the proportion of water storage tissue was larger compared to the cotyledons while air space volume was less. This correlates with the development of functional and efficient C_4_ photosynthesis. Akhani et al. (Akhani *et al*. 2009) who determined the carbon isotope composition of cotyledons and leaves of *H. mollissima* Bunge, found that cotyledons and leaves have C_3_ and C_4_ type isotope values, respectively. This fits with our observations of typical C_4_ genes (PEPC, BCA, and NAD-ME) being higher expressed in the FLs (Fig. 8(A)).

A C_3_ to C_4_ switching mechanism has been reported for several species in the Salsoloideae (Caroxyleae and Salsoleae tribes) and Suadedoideae subfamilies (Rudov *et al*. 2020). Thus, their germination usually coincides with low-temperature conditions that possibly favor C_3_ photosynthesis in the cotyledons. As these species grow and the temperature rises, C_4_ photosynthesis is encouraged in the leaves (Rudov *et al*. 2020). Plants expressing such C_3_ to C_4_ switching may be appropriate models to explore the C_4_ pathway and its regulatory mechanisms due their lack of phylogenetic noises and interspecies differences. Transcriptome analysis of *Haloxylon ammodendron* (Y. Li *et al*. 2015) and *Salsola soda* (Lauterbach, Billakurthi et al. 2017) both belonging to the *Salsoleae* tribe and performing a NADP-ME subtype of C_4_ photosynthesis in true leaves, revealed differentially expressed C_4_ pathway genes, such as PEPC between C_3_ cotyledons and C_4_ true leaves, but paid less attention to possible differences in regulatory pathways. In the present study we investigated potential shoot-root communications during the C_3_ to NAD-ME subtype of C_4_ transition by focusing on the DEGs in hypocotyls before and after the C_4_ first leave’s formation.

### Novel Consistent Data Provided by *De novo* Assembly of *H. mollissima* Bunge Transcriptome

Most transcriptome data analyses in the field of C_4_ photosynthesis, with a few exceptions, have used a reference-based approach to annotate their data using the Arabidopsis data (Gowik and Westhoff 2011; Lauterbach, Billakurthi *et al*. 2017; Lauterbach, Schmidt *et al*. 2017; Siadjeu, Lauterbach and Kadereit 2021). As a non-model species from the Caroxyleae tribe whose genome is not sequenced, we used RNA-seq to de novo assemble and characterize the *H. mollissima* Bunge hypocotyls and FLs transcriptome.

PCA (Fig. 3) and analysis of functional classes of annotated genes (Fig. 4(a)) confirm the sequencing data’s reasonable and comparable relationship. Based on functional annotation, FLs expressed the NAD-ME subtype of the C_4_ photosynthesis pathway. The expression of photosynthesis-related genes was reduced in hypocotyls after establishing the photosynthetic leaves. The formation of FLs also coincides with increased translation processes in the hypocotyls an example of this being the upregulation of the ubiquitin-proteasome system, a critical regulator of many plants signaling pathways in respond to internal and external cues (Sadanandom *et al*. 2012).

### Possible Regulatory Mechanisms Implicated in the Unique C_4_ Features through Adjustment of Shoot and Root Communication

Several regulatory systems with overlapping functions control the developmental processes in plants. These include among others phytohormones, signaling peptides, and complex network of TFs. Long-range regulators are an essential part of these systems. C_4_ photosynthesis which is seen as a response to a reduced atmospheric CO_2_, requires a coordination between the root and the modified shoot. Similarly, the onset of the C_4_ pathway in leaves, followed by increased photosynthesis rate and more efficient use of resources, requires coordination between roots and the modified shoots via signaling molecules. Interspecies grafting experiments between C_3_ *Flaveria robusta* and C_4_ *Flaveria bidentis* indicated that the root preliminary controls sulfur allocation between roots and shoots and sulfur homeostasis in C_4_ *Flaveria* (Gerlich *et al*. 2018). The communication ways between above- and below-ground plant parts are very diverse. They can be physical (e.g. propagating Ca^2+^ waves), chemical (e.g. signaling peptides and hormones), or molecular (e.g. RNA and proteins) (Shabala *et al*. 2015). These signals can move through vasculature systems or cell-by-cell via stems or stem-like tissue (Bartusch and Melnyk 2020). Also, the phloem comprises several mobile developmental signals (mRNAs, proteins, peptides, e.g.), which can interplay in the different stages of development and growth (Koenig and Hoffmann-Benning 2020). Long-range movement of regulatory molecules has already been demonstrated, for auxin (Bellstaedt *et al*. 2019), cytokinin, abscisic acid (Kiba *et al*. 2011), various signaling peptides such as CEP, specific transporters or receptors and some TFs (Shabala *et al*. 2015). By comparing transcriptome profiles of hypocotyls during the C_3_ to C_4_ transition, we revealed that transcripts of genes encoding components of different phytohormone metabolisms, signaling peptides, and TFs (Fig. 6, Additional file 8) were differentially expressed in hypocotyls after establishing the C_4_ leaves. In addition, the expression levels of several transporters, receptors, and responsive genes of root- or leaf-derived signals were also affected, including those of PIN1, CEPR, and ARF7/ARF19 (Fig. 6, Additional file 8). Some of these regulators are most likely implicated in the normal development of hypocotyls. Therefore, to distinguish true C_4_-related genes, we used a literature survey to exclude those genes involved in the hypocotyl development (Additional file 6, Additional file 7). The mobility of mRNA as long-distance signal is supposed to be a complex process while elucidating its physiological roles remains challenging (Xia *et al*. 2018).

### Nitrogen Use Efficiency: The Reduction of Shoot-to-Root Nitrogen Status Signals after the Formation of the C_4_ FLs

Nitrogen assimilation by roots can be controlled by a number of signals representing the nitrogen status in the shoots (Ruffel *et al*. 2008; Forde 2002). These signals include phytohormones such as auxin and cytokinin (Kiba *et al*. 2011). Our results showed that several genes involved in the biosynthesis, transport, and biodegradation of phytohormones were differentially expressed in the hypocotyls after formation of the FLs. Cytokinin signaling increased after the establishment of C_4_ photosynthesis. Shoot-derived cytokinin is a signal of nitrogen satiety in well-supplied nitrogen conditions, suppressing the nitrogen uptake by roots (Kiba *et al*. 2011). Our findings suggest that long-range cytokinin signaling can be a candidate pathway for controlling nitrogen uptake in C_4_ plants. No particular cytokinin transporters for this shoot-to-root movement through the phloem has been identified yet, but ENTs are likely involved (Liu, Zhao and Zhang 2019). Type-B ARR is the positive regulator of cytokinin signaling (Argyros *et al*. 2008) and type-A ARR is the negative regulator of cytokinin signaling (To *et al*. 2004) which have their corresponding transcripts (AT1G67710 and AT3G57040) up- and down-regulated, respectively, in the hypocotyl B.

Due to its role in lateral root initiation, auxin has long been known as a shoot-to-root nitrogen signal (Forde 2002; Fukaki and Tasaka 2009). In the C_4_ crop maize, auxin concentration in phloem exudates decreases under high nitrate concentrations (Tian *et al*. 2008). In the same line, our data showed that cell-to-cell auxin transporters, responsible for auxin’s long-distance movement and an enzyme catalyzing the rate-limiting step in auxin biosynthesis, were down-regulated in hypocotyls B. In contrast, the intracellular auxin transporters that modulate the nuclear abundance of auxin by transporting auxin into the ER lumen (Sauer and Kleine-Vehn 2019) were up-regulated. These findings imply that long-distance movement of auxin is reduced in hypocotyls after C_4_ leaf formation. At the same time, the NRT1.1 transporter becomes down-regulated in hypocotyls B. As a dual-affinity transporter, NRT1.1 acts as a nitrate sensor that regulates the expression level of genes involved in nitrate transport (Sun and Zheng 2015). It is located in roots, vascular cells, and shoots and displays an auxin-responsive activity. The reduced expression in hypocotyl B is consistent with other findings in this study, implying a general reduction of auxin content, either derived from leaves or the hypocotyl itself, in the hypocotyl B. Furthermore, NRT1.1 has a significant role in the nitrogen uptake in roots affected by shoot-derived signals. Thus, its lower expression in roots after formation of the first leaves (Fig. 8(c)) may be a response to a decrease in the nitrogen uptake by roots after C_4_ establishment. Other reports claim, however, that local increases of auxin biosynthesis and transport in developing leaves is necessary for manifestation of C_4_ key property of higher vein density (Huang *et al*. 2017).

A further mobile regulator mediating nitrogen assimilation in roots under nitrogen deficiency condition, is CEP which has a shoot orientation from the roots. CEP receptor, like CEPR2, was down-regulated in hypocotyl B (Fig. 8(d)), most likely due to the efficient use of nitrogen by C_4_ plants. This likely reflects the reduced effect of root-derived CEP signal after C_4_ leave’s formation. Indeed, this receptor implicates nitrogen uptake signaling by enhancing the expression of NRT1(Tabata *et al*. 2014; Ohkubo *et al*. 2017). The observed lower NRT1 expression in roots after C_4_ first leave’s formation (Fig. 8(c)) is thus consistent with possible reduced root-derived CEP. Notably, CEPR2 expression was higher in the cotyledon B (C_3_) compared to the FLs (C_4_) (Fig. 8(a)), indicating a higher level of nitrogen consumption by C_3_ cotyledons than by C_4_ leaves.

Taken together, gene expression changes after emerging C_4_ photosynthesis in the FLs indicate modulated auxin and cytokinin metabolism and signaling alongside a reduced CEP signaling in hypocotyl B and FLs. This supports the notion of an enhanced nitrogen use efficiency and reduction in total nitrogen requirement in C_4_ plants (Jobe *et al*. 2020).

### Drought Resistance and Water Use Efficiency: Several Pathways involved in the High-Temperature Resistance after the Formation of the C_4_ FLs

Through different mechanisms, C_4_ plants produce more efficiently than C_3_ under higher temperatures. One mechanism is preventing dehydration through stomatal closure. Stomatal closure under drought stress is regulated by root-derived abscisic acid (Kuromori, Seo and Shinozaki 2018). Some studies claim that stomata of C_4_ plants are more sensitive to intercellular CO_2_ concentration (Ci) compared to C_3_ (Huxman and Monson 2003), and that abscisic acid can increase this sensitivity (Dubbe, Farquhar and Raschke 1978). We observed that the expression level of several abscisic acid influx genes (ABCG30, ABCG40) (Zhang *et al*. 2014; Kang *et al*. 2015) was up-regulated in hypocotyls B compared to hypocotyls A (Fig. 6, Fig. 8(d)). Our finding for up-regulation of ABCG40 in C_4_ leaves compared to C_3_ cotyledons (Fig. 8(a)) is consistent with its functions in shoots as a root-derived-abscisic acid importer and its essential role in stomatal closure (Kang *et al*. 2010). The differential regulation of ABCG40 in roots (Fig. 8(d)) likely is due the interaction between abscisic acid and nitrate uptake (Harris and Ondzighi-Assoume 2017). Moreover, studies on the amphibious *Eleocharis vivipara* have revealed the significant role of abscisic acid and hormonal control in the manifestation of Kranz anatomy (Ueno 1998).

### Stress- and Light-Responsive Differentially expressed Mobile TFs in Hypocotyls are Possibly Involved in the Formation of C_4_ Unique Features

Many of the TFs that are up-regulated in the hypocotyls after the formation of the FLs are implicated in the regulatory networks controlling responses to the different plant development and growth conditions as well as abiotic stresses. The appearance of C_4_ leaves in *H. mollissima* Bunge, and other halophytes or xerohalophytes switching chenopods is coincides with an increase in environmental temperature (Rudov *et al*. 2020). Transcriptome analysis of C_3_ cotyledons and C_4_ leaves in *Salsola soda* demonstrated that 55 and 32 TFs were increased and decreased in C_4_ leaves compared to C_3_ cotyledons, respectively (Lauterbach, Billakurthi et al. 2017). Not surprisingly, a greater number of the up-regulated TFs in our study were stress-responsive, including NAC (Shao, Wang and Tang, 2015), WRKY (Wang *et al*. 2018), and NF-Y (Zhao *et al*. 2017).

Among the differentially expressed TFs in hypocotyls before and after the formation of the C_4_ FLs, SHR and HY5 TFs are known to be mobile although at protein level (Dolan 2001; Chen *et al*. 2016). SHR is implicated in modulating endodermis (known as bundle-sheath) differentiation in leaves and apparently plays a role in the Kranz anatomy formation in maize (Slewinski *et al*. 2012). Moreover, Siadjeu et al. (Siadjeu, Lauterbach and Kadereit 2021) who compared transcriptome data of C_3_, C_2_, and C_4_ species from *Camphorosmeae*, found that, BBX15, SHR, SCZ, and LBD41 are more expressed and co-regulated in the C_4_ than C_3_ species. Interestingly, out of 38 differentially expressed Arabidopsis orthologous TFs in the comparative transcriptomic analysis between Kranz (foliar leaf blade) and non-Kranz (husk leaf sheath) maize leaves, we identified four TFs that express more in hypocotyl after C_4_ leave’s formation; including SHR (AT4G37650), BHLH96 (AT1G72210), GATA7 (AT4G36240) and WRKY12 (AT2G44745).

Most of the up-regulated TFs after C_4_ manifestation belonged to C2H2 and bZIP families, including HY5. HY5 is a master regulator of transcription (Gangappa and Botto 2016), a phloem-mediated shoot-to-root signal for lateral root formation, and thus the carbon-nitrogen balance adjustment in Arabidopsis (Burko *et al*. 2020). HY5 mediates photomorphogenesis so that it is located downstream of phytochrome B, the sensor of light and temperature (Legris *et al*. 2016). Thus, in C_4_ plants where shoots are exposed to high light intensity and temperature, and since the function of HY5 is conserved in many plants (Yamawaki *et al*. 2011; Huai, Jing and Lin 2020), HY5 can be an appropriate candidate for synchronizing shoots and roots. Moreover, HY5 is one of the light-responsive TFs essential in inhibiting hypocotyl elongation during seedling (Oyama, Shimura and Okada 1997). Therefore, HY5 is a shared regulator between several various pathways.

The mRNA of 12 differentially expressed TFs in the *H. mollissima* Bunge hypocotyls have cell-to-cell mobility, according to the TAIR database (Table 1). We also predicted the probable cis-elements present in the promoter region of their orthologous in *A. thaliana*. Several of the predicted cis-elements in the mobile TFs were light- and stress-responsive (Additional file 10). This indicates that expression and transport of these TFs is influenced by abiotic stresses. One of these TFs is WRKY40, an orthologue of which, ZmWRKY40, is a drought-responsive gene in *Zea mays* (Wang *et al*. 2018). HSFA1e was more expressed in hypocotyl B, which is implicated in the tolerance to heat shock stress via transcriptional regulation of HsfA2 function (Nishizawa-Yokoi *et al*. 2011). Overexpression of maize ZmHsf06 in Arabidopsis, improves drought-stress tolerance (H. Li *et al*. 2015). The findings of this study pave the way for identifying possible regulators underlying the coordination of developmental changes associated with functional C_4_ photosynthesis. Also, by investigating a plant with a NAD-ME subtype of C_4_ photosynthesis our data fills the gap caused by previous studies solely concentrating on the NADP-ME subtype of photosynthesis. Our data provides a valuable source for further functional genomics and genetic studies to identify master regulators of C_4_-related traits and engineering C_4_ features into C_3_ crops.

## Abbreviations

TF: Transcription factor
DEG: Differentially expressed gene
FLs: First leaves
PEPC: Phosphoenolpyruvate carboxylase
BCA: Beta-carbonic anhydrase
PPdK: pyruvate orthophosphate dikinase
Asp-AT: Aspartate aminotransferas
Ala-AT: Alanine aminotransferase
NAD-ME: NAD-dependent malic enzyme
NADP-ME: NADP-dependent malic enzyme
TPT: Triosephosphate translocator
PPase6: Pyrophosphatase
PPdK-RP1: Pyruvate orthophosphate dikinase related protein1
PPT2: PEP/phosphate translocator
BASS4: Bile acid:sodium symporter family protein
BASS2: Bile acid:sodium symporter family protein
AMK2: Adenosine monophosphate kinase
HY5: Elongated hypocotyl 5
SHR: Short-Root

## Author Contribution

MZ performed the experiments, analyzed data, and wrote the original draft. TR performed the microscopic analysis and critically revised the manuscript. MRG analyzed data. AMB-M conceived and designed the research, supervised the experiment, and critically revised the manuscript. All authors contributed to the article and approved the submitted version.

## Acknowledgment

We thank Stanislav Kopriva (University of Cologne) for the critical reading of the manuscript. We also would like to thank Alexander Rudov and Hossein Akhani (University of Tehran) for their support during seed collection.

## Additional information

### Supplementary Information

**Supplementary file 1** Summary of A) sequencing information B) De novo transcriptome assembly results by Trinity

**Supplementary file 2**. The accession number of uploaded raw sequences to ??

**Supplementary file 3**. Active functional classes in the First leaves, hypocotyl A and hypocotyl B

**Supplementary file 4**. Abundance (total mean TPM) of C_4_-related proteins and genes categorized in the CRP and NCRP signaling peptide groups

**Supplementary file 5**. Transcript abundance, statistics of differentially expressed genes, and their annotations.

**Additional file 6**. Venn diagram analysis of DEGs of two developmental *H.mollissima*’s hypocotyls and identified genes related to the hypocotyl developments in flax (Roach and Deyholos, 2008) and hemp (Behr *et al*. 2018) based on their assigned Arabidopsis ID

**Supplementary file 7**. List of the unique and shared genes compared to DEGs of two developmental *H.mollissima*’s hypocotyls and identified genes related to the hypocotyl developments in flax (Roach and Deyholos, 2008) and hemp (Behr *et al*. 2018) based on their assigned Arabidopsis ID.

**Supplementary file 8**. Functional analysis of the Cell signaling genes using Mapman

**Supplementary file 9**. Differentially transcription factor genes between hypocotyl A and hypocotyl B using PlantTFDB v5.0

**Supplementary file 10**. Cis-elements located upstream of orthologous transcription factor genes with mobile mRNA in Arabidopsis

**Additional file 11** Validation of RNA-Seq results using quantitative RT-PCR (qRT-PCR)

