## Additional File 6 @ 11 for "Comparative Transcriptome Analysis of Hypocotyls During the Developmental Transition of C_3_ cotyledons to C_4_ Leaves in *Halimocnemis mollissima* Bunge"

**Additional file 6** Venn diagram analysis of DEGs of two developmental *H.mollissima*'s hypocotyls and identified genes related to the hypocotyl developments

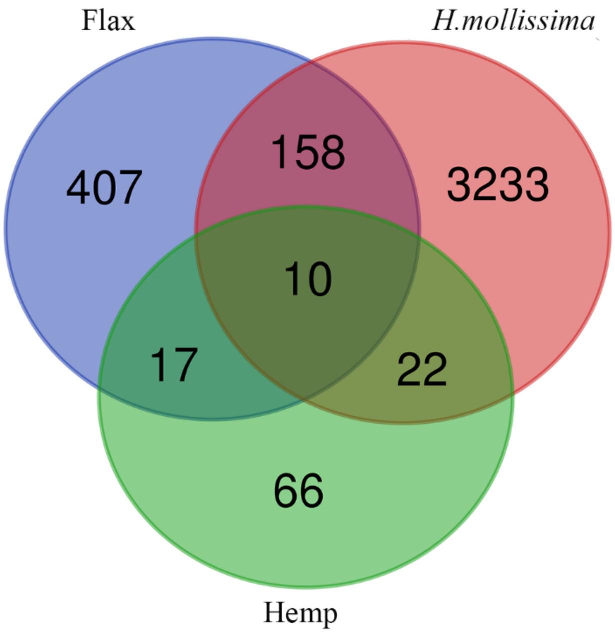


**Additional file 11** Validation of RNA-Seq results using quantitative RT-PCR (qRT-PCR)

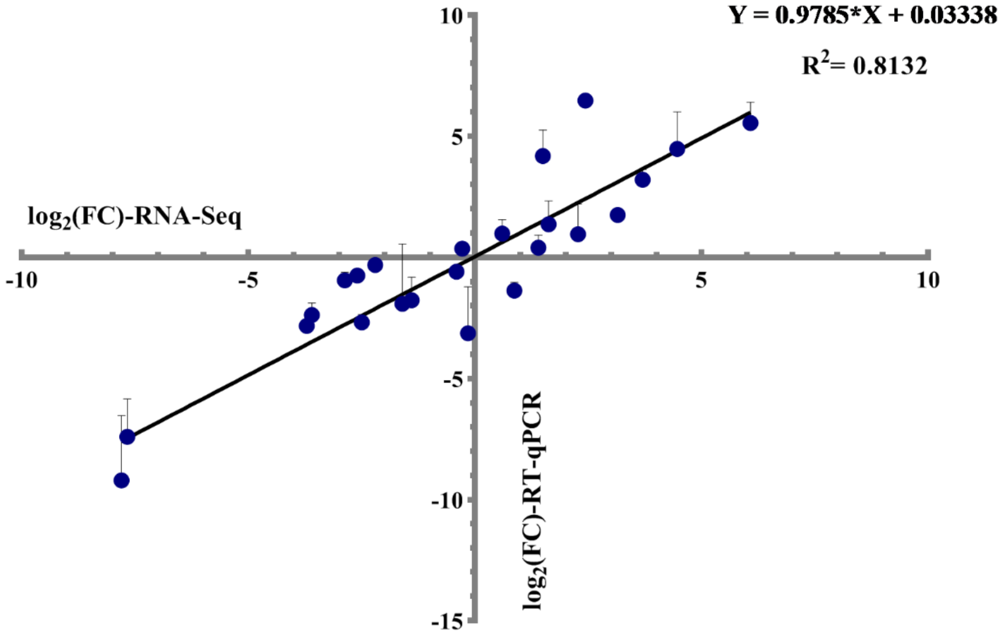
