## Additional File 1 for "Comparative Transcriptome Analysis of Hypocotyls During the Developmental Transition of C_3_ cotyledons to C_4_ Leaves in *Halimocnemis mollissima* Bunge"

**Supplementary file 1**: summery of A) Sequencing information B) De novo transcriptome assembly results by Trinity

| **A) Summary of sequencing information** | | | | | | |
|---|---|---|---|---|---|---|
| **Replicates** | Fist leaves (1) | First leaves (2) | Hypocotyl A (1) | Hypocotyl A (2) | Hypocotyl B (1) | Hypocotyl B (2) |
| **Number of row reads** | 20,180,458 | 19,230,133 | 19,825,621 | 20,536,077 | 20,702,522 | 161,868,549 |
| **Sequence length** | 150 | 150 | 150 | 150 | 150 | 150 |

| **B) Statistics of de novo transcriptome assembly using Trinity** | |
|---|---|
| **Number of contigs (>= 0 bp)** | 280973 |
| **Number of contigs (>= 1000 bp)** | 74510 |
| **Largest contig** | 17530 |
| **Total length (>= 0 bp)** | 256,275,034 |
| **Total length (>= 1000 bp)** | 173,811,615 |
| **N50** | 1847 |
