## Additional File 2 for "Comparative Transcriptome Analysis of Hypocotyls During the Developmental Transition of C_3_ cotyledons to C_4_ Leaves in *Halimocnemis mollissima* Bunge"

Supplementary file 2: The accession number of uploaded raw sequences to …

| Samples (Tissues) | ID | Accession Number |
|---|---|---|
| Fist leaves | FL_1 | |
| | FL_2 | |
| Hypocotyls A | H_A1 | |
| | H_A2 | |
| Hypocotyl B | H_B1 | |
| | H_B2 | |
